# Multiscale transcriptomic organization of the human brain with DigitalBrain

**DOI:** 10.64898/2026.04.14.718492

**Authors:** Jiangliang An, Xiangyang Hu, Yunhan Jiang, Mingyu Jiang, Shuowen Qiu, Guole Liu, Xunbin Wei, Yilong Wang, Julie Qiaojin Lin, Cunyu Wang, Meng Lu

**Author notes:** Correspondence to: Meng Lu. These authors contributed equally to this work (Jiangliang An, Xiangyang Hu).

## Abstract

The human brain varies across anatomical regions, cell types, development, aging and disease states, yet existing single-cell transcriptomic resources remain fragmented and difficult to integrate into a unified biological model. Here we present DigitalBrain, a human brain-specific atlas and foundation-model framework for organizing diverse and fragmented human brain transcriptomic data across scales. We first built DigitalBrain-Atlas, a harmonized whole-brain single-cell resource comprising 16.35 million transcriptomes from 2,143 donors across 165 brain regions, spanning the human lifespan and multiple neurological and clinical conditions. We then developed DigitalBrain-M1, a Transformer-based model that jointly encodes gene identity and expression magnitude to learn a shared embedding space for cells and genes. Across held-out datasets, DigitalBrain supported robust single-cell integration, clustering and cell-type annotation while preserving major biological structure and reducing technical fragmentation. Beyond these benchmarks, the learned embeddings revealed emergent large-scale hierarchical organization of the human brain, linking anatomically distinct regions into higher-order patterns consistent with known functional systems. Applied to human hippocampal aging, DigitalBrain identified cell-type-specific aging sensitive gene sets, highlighted dentate gyrus granule cells as a particularly age-sensitive population, and discovered selective reorganization of gene programs related to synaptic transmission, postsynaptic structure, membrane excitability and axon guidance during aging. Cross-dataset convergence was strongest at the level of functional modules and recurrent aging sensitive genes. Together, these results demonstrate that DigitalBrain is a brain-specific framework for mapping human brain organization across scales, and as an early step towards a complete virtual organ for the human brain.

## Introduction

The human brain is organized across multiple, interdependent axes of variation, including anatomical region, cell identity, developmental history, aging and disease state^1^. Single-cell transcriptomics has greatly expanded our ability to catalogue this diversity^2,3^, but a unified modelling framework that learns coupled representations of cells and genes is still lacking^4^. Such a framework would require not only a harmonized whole-brain atlas spanning regions, life stages and disease conditions, but also a brain-specific representation model capable of learning a unified embedding space for cells and genes, thereby enabling integration, annotation and discovery across the full breadth of existing data.

Existing human brain single-cell datasets remain distributed across studies focused on specific regions, cell populations, developmental stages and disease contexts^5–8^. As a result, the full spectrum of human brain cellular diversity remains only partially sampled across brain structures and life stages. Although recent efforts such as Siletti et al. (2023)^9^ have provided whole-brain datasets covering major anatomical regions, these datasets remain limited in donor number and primarily capture a narrow segment of the lifespan. More recently, Chen et al. (2024)^10^ expanded this scale to 11.3 million human cells and nuclei by integrating 70 studies. While this atlas represents a major advance toward cross-region integration, residual heterogeneity in source level anatomical annotations and metadata granularity may still complicate systematic comparisons across integrated datasets. Importantly, however, the challenge is not simply one of scale: datasets remain highly heterogeneous in anatomical nomenclature, sampling strategy, cell type annotation, disease context and quality control, making cross-study integration difficult and obscuring shared biological structure^4,11^.

We therefore reasoned that a useful brain-specific representation model would require more than a large training corpus; it would require a harmonized whole-brain atlas that defines the observable biological state space of the human brain across regions, cell populations, life stages and disease conditions. Such an atlas would need to integrate not only fragmented datasets across brain regions but also the fragmented landscape of curation practices, establishing a unified space for representing human brain biology. In this view, the atlas is not only a resource for description, but also the substrate for learning a unified representation: one that can support advanced, large-scale analyses and, crucially, serve as a foundation for constructing AI models capable of capturing the brain’s hierarchical organization, cell-type and cell-state dynamics, and principles of gene regulation.

Here we present DigitalBrain, a resource-grounded AI framework built on this principle. We first collected 109 human brain datasets and assembled a curated whole-brain single-cell atlas comprising more than 16 million cells from 2143 human donors, spanning 165 brain regions, full human lifespan and multiple neurological conditions. Using this harmonized atlas, we trained a human brain-specific model, DigitalBrain-M1, to learn coupled representations of cells and genes within a unified modelling framework.

We show that DigitalBrain learns biologically meaningful representations across datasets, cell populations and molecular programs. At the cell level, the model supports cross-dataset integration, cell type annotation transfer and preservation of continuous state-associated variation across aging. At the gene level, the learned embeddings recapitulate biologically coherent functional modules and provide a basis for measuring the of the gene functional change at different cellular context. As a proof-of-principle application, we applied this framework to human hippocampal aging, and identified dentate gyrus granule cells showing a strong aging-associated shift in embedding space, and found that age-variable gene programs were enriched for metabolism, energy regulation and homeostatic maintenance, whereas comparatively stable programs were enriched for neuronal identity and synaptic organization, consistent with the broader view that aging involves selective reorganization of gene–gene coordination rather than uniform collapse^12^. Together, these results establish DigitalBrain as a brain-specific resource and AI platform for organizing human brain diversity and generating interpretable biological hypotheses across physiological and disease states.

## Results

### DigitalBrain-Atlas

Developing a high-performance foundation model of the human brain requires more than scale alone; it relies on high-quality corpus with standardized biological and anatomical structure for robust training and downstream analysis. To address this, we first constructed DigitalBrain-Atlas, a harmonized single-cell resource comprising 16,351,058 transcriptomes from 2143 donors across (109 datasets) (Fig. 1a-c). The atlas spans 165 brain regions, the human lifespan from 0 to 96 years (Fig. 1d) and a great variety of cell type (Fig. 1e). We organized these data within a standardized multi-level anatomical framework based on the Allen Adult Human Brain Atlas^13^, enabling each cell to be mapped into a common hierarchical representation of brain structure (Fig. 1f) and disease type (Fig. 1g). In parallel, we reconciled study-specific metadata and cell annotations through a curated harmonization pipeline that standardized region labels, donor/sample metadata, disease labels, and major cell-type assignments across studies (Fig. 1h).

**Fig. 1.**
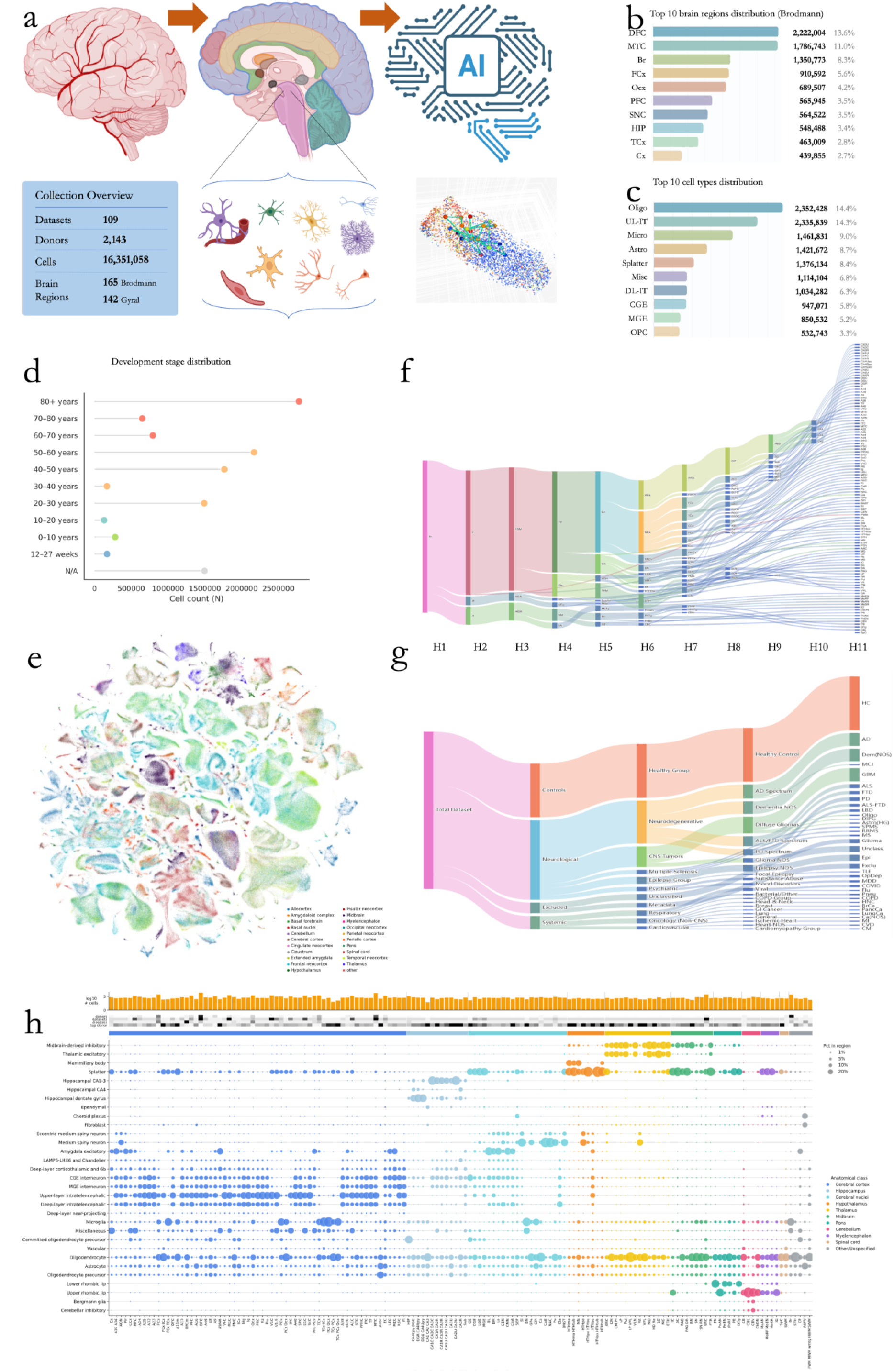
Overview of the DigitalBrain atlas, its composition, and anatomical coverage. **(a)** Summary schematic and collection statistics for the aggregated dataset, including the numbers of datasets, donors, cells, and harmonized brain regions. **(b)** Cell-count distribution across the 10 most highly represented brain regions, annotated using the modified Brodmann framework. **(c)** Cell-count distribution across the 10 most abundant major cell types in the atlas. **(d)** Distribution of cells across developmental stages and age groups represented in the dataset. **(e)** UMAP visualization of a subset of atlas cells, colored by main brain structures. **(f)** Sankey diagram showing coverage across the 11-level anatomical hierarchy used in DigitalBrain; node sizes and link widths are scaled by cell abundance (log). **(g)** Sankey diagram summarizing disease and condition coverage across the integrated dataset. **(h)** Anatomy-guided bubble map showing cell-type composition across human brain regions ordered according to the Allen Brain Atlas. Bubble size indicates the fraction of each cell type within each region, and colors follow the major anatomical class scheme. Annotation tracks above the panel summarize regional cell number, donor coverage, contributing datasets, disease-state coverage, and donor dominance.

To assess the breadth and cellular composition of the assembled DigitalBrain-Atlas, we examined donor-level cell-class composition across all integrated resources. Figure Table 1 visualizes each donor as a vertical bar, with stacked colors representing the proportions of neuronal, glial, and non-parenchymal populations. Donors are grouped first by sample system (e.g., primary adult brain, developmental tissue, organoid models, tumour-associated samples, and other specialized CNS datasets) and then by individual collection; white gaps denote collection boundaries. The upper annotation tracks indicate sample system and collection identity, while the lower grayscale track encodes donor age stage, spanning from prenatal to aged. This overview highlights substantial inter-donor variation in cellular composition both within and across collections, underscoring the heterogeneity captured by the atlas and the importance of harmonized integration for downstream analysis.

**Table 1.**
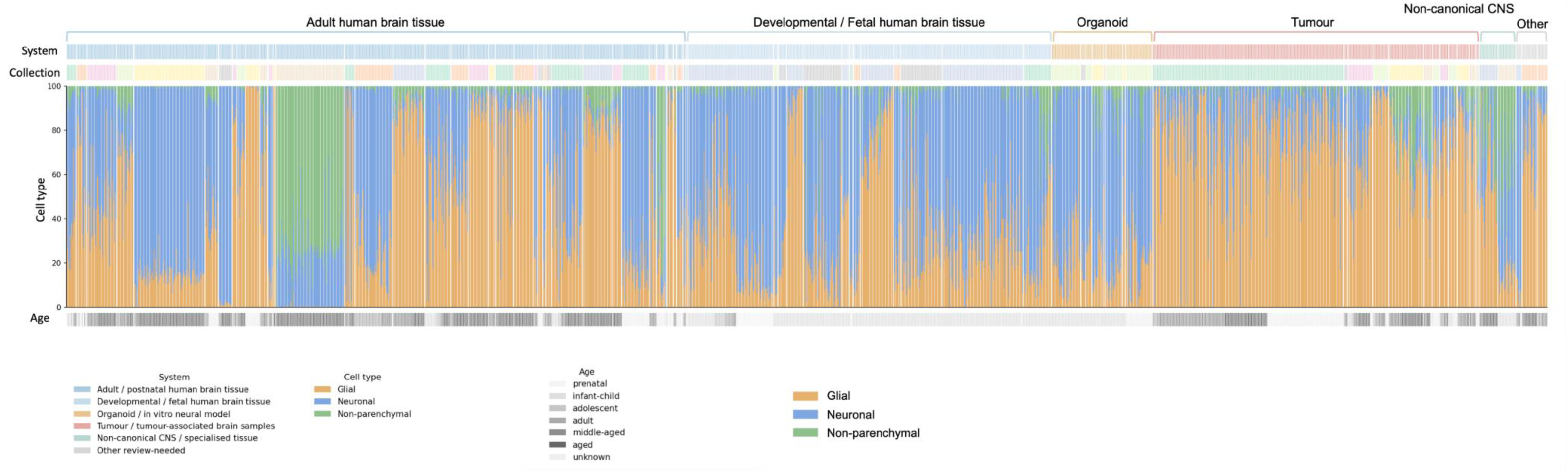
Donor-level cell-class composition across integrated human brain resources. Each vertical bar represents one donor aggregated within collection. Stacked colors indicate the proportion of cells assigned to the three major cell classes, including neuronal, glial and non-parenchymal populations. Donors are grouped first by sample system and then by collection, with white gaps indicating collection boundaries. The upper annotation tracks denote sample system and collection identity, and the lower grayscale track indicates donor age stage, ranging from prenatal to aged. This view summarizes inter-donor variation in cellular composition across primary adult and developmental brain tissue, organoid models, tumour-associated samples and other specialised central nervous system datasets.

The resulting atlas contains neurons, glial populations, vascular cells, fibroblasts, and related non-neuronal cell classes, with broad representation across cortical and subcortical systems. Together, these features make DigitalBrain-Atlas a unified training corpus for learning brain-specific representations across datasets rather than a simple aggregation of disconnected studies. An interactive version of the atlas, together with open access to the curated dataset for download, is available through the DigitalBrain Data Explorer (https://lumenglab.github.io/digitalbrain-data-explorer/).

### A hierarchical single-cell foundation model of the human brain

We next developed DigitalBrain-M1, a brain-specific Transformer model trained on DigitalBrain-Atlas to learn embedding space for cells and genes across the human brain. Unlike foundation models trained on relatively flat collections of dissociated cells^14–19^, DigitalBrain-M1 was built on a hierarchically organized atlas spanning whole-brain anatomy, 11 levels of anatomical organization, regional identity and single-cell transcriptomic state. This whole brain atlas training enables the model to learn not only cell-intrinsic transcriptional programs, but also patterns embedded within the multiscale organization of the brain. In this sense, DigitalBrain provides a virtualized representation of the human brain at single-cell resolution, while remaining a transcriptomic foundation model rather than a mechanistic simulation of organ function.

The model takes single cell specific ranked gene-expression profiles as input and encodes each gene token using two complementary features: a gene identity embedding, which encodes functional identity, and an expression-value embedding, which captures quantitative abundance. For each cell, the top 2,048 non-zero highly expressed genes of each cell are encoded as the primary sequence^18^, enabling the model to focus on the dominant transcriptional features that define cellular identity (Fig. 2a).

**Fig. 2.**
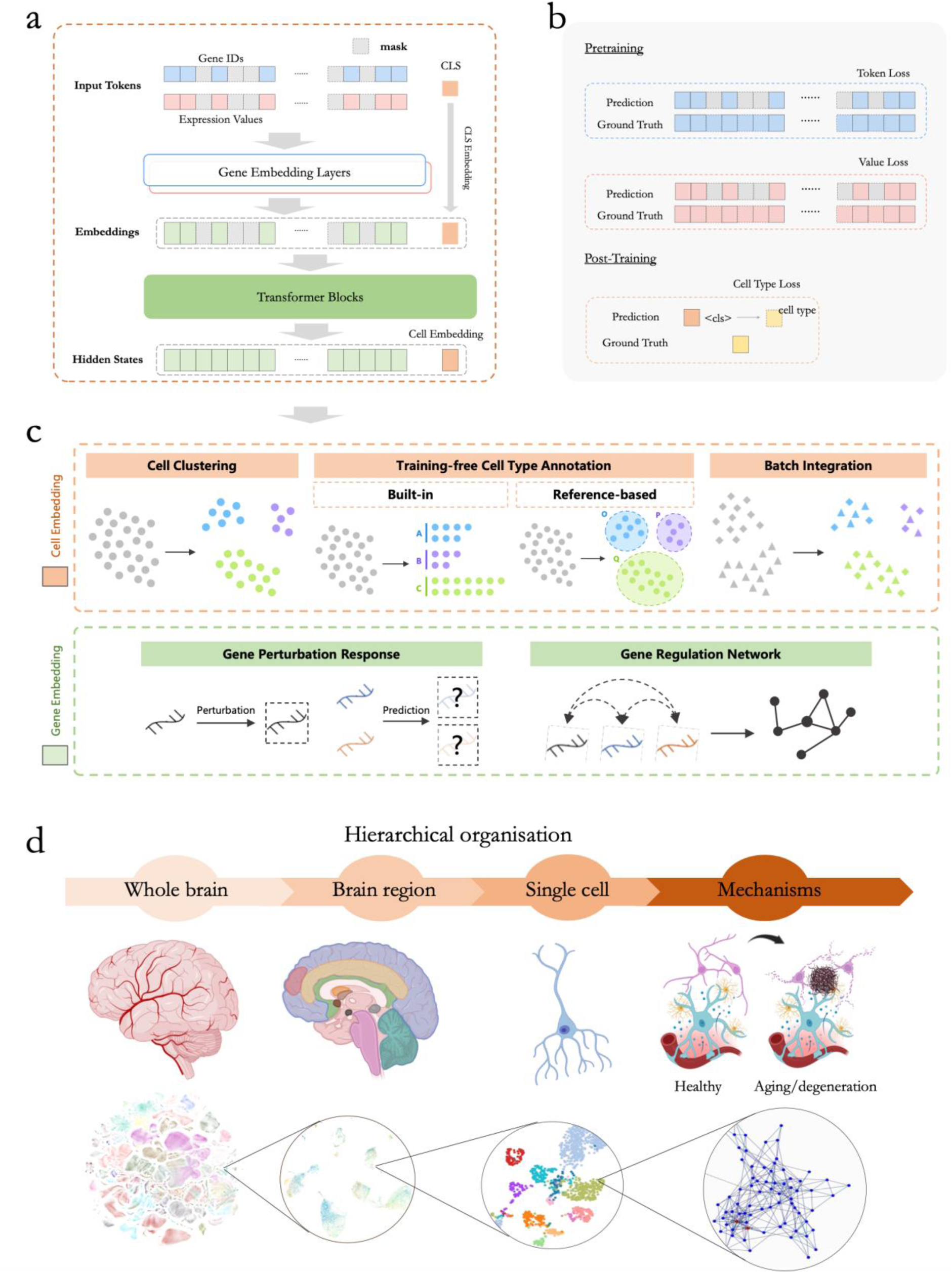
The DigitalBrain framework: model architecture, training strategy, and downstream applications. **(a)** Schematic of the DigitalBrain input architecture. For each cell, the top 2,048 expressed non-zero genes are encoded as the primary input sequence. Each token combines a gene identity embedding and an expression-value embedding, and a [CLS] token is prepended to capture the global cell state. **(b)** Training strategy for DigitalBrain. During self-supervised pretraining, the model is optimized using masked gene identity prediction and masked expression-value prediction. During post-training, the cls embedding is further optimized with supervised cell-type classification loss, adapting the representation to brain-specific cellular identities. **(c)** Overview of downstream applications enabled by the learned representation space. Cell embeddings support cell clustering, training-free cell-type annotation using either built-in labels or reference-based transfer, and batch integration across datasets. Gene embeddings support analyses of perturbation responses and inference of gene regulatory relationships from learned gene–gene structure. **(d)** Conceptual illustration of the hierarchical biological scales captured by DigitalBrain, spanning whole brain, brain region, single cell, and molecular mechanisms, including healthy and ageing/degenerative states.

Training proceeded in two stages (Fig. 2b). We first performed self-supervised pretraining using masked gene identity prediction and masked expression prediction, enabling the model to learn contextual dependencies among genes directly from the atlas. We then carried out brain-specific post-training on 12 curated human brain datasets using 31 core cell types standardized to the Human Brain Cell Atlas^9^. This second stage aligns the learned embedding space with established brain cell identities and adapts the model to the cellular structure of the human brain.

DigitalBrain-M1 produces a shared representation space that supports analyses at both the cell and gene levels (Fig. 2c). At the cell level, the learned embeddings can be used for integration, clustering, and reference-based annotation across datasets. At the gene level, context-specific embeddings can be extracted from defined cell types, brain regions, or phenotypic states, providing a basis for analyzing gene programs and context-dependent molecular relationships. Together, these representations enable analyses across multiple biological scales, from individual cells to brain subregions (Fig. 2d), and provide the foundation for the downstream applications described below.

### Biologically meaningful representations at both the cell and gene levels

To evaluate whether DigitalBrain-M1 learns a biologically meaningful representation, we benchmarked the model at both the cell and gene levels. At the cell level, we assessed integration, clustering, and cell-type annotation across heterogeneous single-cell brain datasets. At the gene level, we asked whether the learned embeddings capture structured functional relationships, including cell-population-specific gene programs and known biological pathways.

At the cellular level, the learned embedding preserved major biological structure. In a UMAP projection of the curated atlas subset, major cell classes formed distinct groups, consistent with separation by biological identity rather than collapse of diverse populations into a single latent space (Fig. 3a). When applied to 12 held-out test datasets, DigitalBrain aligned cells across studies and platforms while maintaining separation of major biological classes, supporting generalization to unseen data (Fig. 3b).

**Fig. 3.**
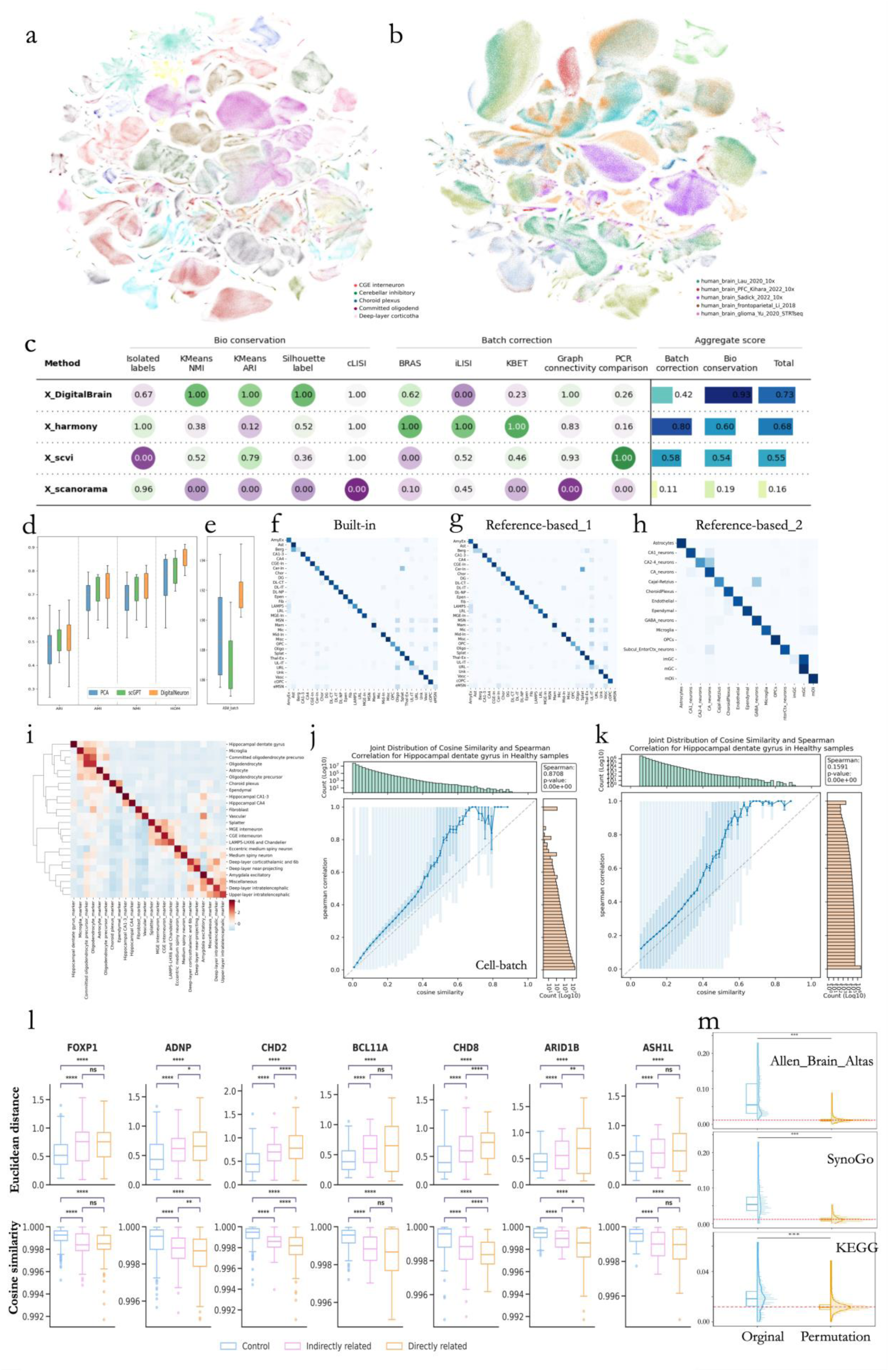
DigitalBrain learns a unified representation space across cell-level integration tasks and gene-level functional structure. **(a)** UMAP of a subset of the curated brain atlas, colored by predicted major cell types. Separation of major populations is consistent with preservation of large-scale biological structure in the learned embedding. **(b)** UMAP of 12 held-out test datasets, colored by dataset source, showing alignment across unseen studies. **(c)** Summary benchmarking with the scIB framework across integration methods, reporting biological conservation, batch correction, and aggregate scores. **(d–e)** Distributions of evaluation metrics across the 12 test datasets for (d) biological conservation (ARI, AMI, NMI, HOM) and (e) batch correction (ASW_batch), compared with baseline methods. **(f–h)** Confusion matrices for cell-type annotation in three settings: (f) end-to-end annotation on independent test datasets, (g) reference-based transfer using cell type annotation in HBCA as the reference, and (H) self-referenced label propagation within the Zhou dataset. **(i)** For each cell type, marker genes were identified from the original single-cell expression data, and their mean pairwise cosine similarity was calculated across gene embedding spaces derived from different cell types. The resulting similarity matrix was row-wise z-score normalized and ordered by hierarchical clustering. Warmer colors indicate greater relative cohesion of a marker set within a given embedding space. **(j-k)** Relationship between gene-embedding cosine similarity and gene-expression Spearman correlation in healthy hippocampal dentate gyrus cells, shown for (j) cell-level (“Cell-batch”) and **(k)** donor-aggregated pseudobulk (“Pseudo-batch”) analyses. Blue lines indicate binned means; error bars denote SEM; shaded regions indicate the full range and interquartile range within bins; marginal histograms show the distributions of each variable. **(l)** Quantitative assessment of gene embedding shifts following in silico knockout of ASD high-risk genes. Boxplots show perturbation-induced changes in gene embeddings after knockout of *FOXP1, ADNP, CHD2, BCL11A, CHD8, ARID1B* and *ASH1L*, measured by Euclidean Distance (up) and Cosine Similarity (down) across three gene categories: Control, Indirectly related and Directly related groups. The distributions highlight stronger embedding perturbation in directly related genes and intermediate effects in indirectly related genes relative to background controls. Statistical significance is indicated by asterisks (**** *P* < 0.0001, *** *P* < 0.001, ** *P* < 0.01, * *P* < 0.05, ns: not significant). The center line indicates the median, box limits indicate the upper and lower quartiles, and whiskers represent the 1.5× interquartile range. **(m)** Raincloud plots showing the distribution of mean within-set cosine similarity for annotated biological gene sets (“Original”) compared with size-matched random gene sets (“Permutation”) across Allen Brain Atlas, SynGO, and KEGG databases. The dashed red line indicates the global background similarity of the network; significance was assessed by permutation testing.

Quantitative benchmarking confirmed these trends. Across integration benchmarks, DigitalBrain-M1 performed strongly in the scIB framework^11^, achieving the highest biological fidelity while maintaining effective batch-correction performance compared to scVI^20^, Harmony^21^, and Scanorama^22^ (Fig. 3c). Across the 12 test datasets, the model achieved favorable scores on biological conservation metrics, including ARI, AMI, NMI, and HOM, while maintaining competitive performance in batch mixing as measured by ASW_batch compared to PCA^23^and scGPT^16^ (Fig. 3d,e). These results indicate that the embedding retains biologically informative variation while supporting cross-dataset integration.

DigitalBrain also supported cell-type annotation in multiple settings. In end-to-end annotation on independent test datasets, the model achieved 87.58% accuracy with a macro F1 score of 0.7626 (Fig. 3f). In a transfer setting using HBCA^9^ as the reference, it achieved 85.35% accuracy with a macro F1 score of 0.7094 (Fig. 3g). In self-referenced label propagation within the Zhou dataset^24^, using 10% of cells as reference for the remaining 90%, the model achieved 92.01% accuracy (Fig. 3h). Together, these analyses show that the learned cell embedding space supports integration, annotation transfer, and robust label propagation across datasets.

We next asked whether cell-population-specific gene relationship networks retained known molecular identity. We first examined whether similarity in the learned gene-embedding space reflected transcriptional concordance. In healthy hippocampal dentate gyrus cells, gene-embedding cosine similarity was positively associated with gene-expression Spearman correlation at both the cell level and the donor-aggregated pseudobulk level (Fig. 3I and J). The relationship was strongest in the cell-level analysis and remained detectable after aggregation, indicating that genes positioned closer in the DigitalBrain embedding space tend to show more coordinated expression patterns. Next, enrichment analysis against curated marker gene sets showed a strong diagonal structure, with each population-derived network most strongly associated with its corresponding marker set (Fig. 3k), indicating that the learned gene relationships preserve cell-type-specific transcriptional programs. Related populations also showed selective off-diagonal enrichment, including overlap within the oligodendrocyte lineage, among interneuron subclasses, and across excitatory neuronal groups, consistent with shared lineage or functional similarity. Together, these results support the biological validity of the DigitalBrain gene embedding space and show that it captures both discrete cell identity and higher-order relationships among related cell populations. These results support the biological relevance of the learned gene embeddings and suggest that they capture structured relationships beyond random proximity in latent space.

To further test whether the embedding space reflects known functional organization beyond pairwise gene–gene relationships, we next examined the internal coherence of annotated biological gene sets. Using raincloud plots, we compared the distribution of mean cosine similarity for gene pairs within each annotated gene set (“Original”) against that of size-matched random gene sets (“Permutation”) across the Allen Brain Atlas, SynGO, and KEGG databases from Enrichr^25^ (Fig. 3m). For each set, the within-set score was computed as the average cosine similarity among all gene pairs belonging to the same set, while the background reference represents the average cosine similarity of gene pairs outside the set. Size-matched random gene sets were repeatedly sampled to generate a null distribution, controlling for set size when evaluating whether a pathway exhibits greater internal coherence than expected by chance. Annotated gene sets consistently showed higher internal cosine similarity compared to random sets (dashed red line indicates background similarity level), with statistical significance assessed by permutation testing. Together, these analyses demonstrate that the gene embeddings capture not only cell-type-specific regulatory structure and single-cell transcriptional concordance, but also coherent functional organization at the level of established biological pathways.

To investigate whether DigitalBrain captures higher-order gene regulatory structure beyond simple co-expression, we performed an *in silico* knockout analysis focusing on high-confidence autism spectrum disorder (ASD) risk genes curated from Li et al. (2023)^26^, Crucially, all ASD datasets were entirely withheld from the training corpus. Furthermore, to circumvents the systematic and confounding biases inherent to experimental perturbation screens^27^, we grounded our predictions in observational cortical transcriptomic data from human ASD donors^28^. Rather than simply tracking expression level changes, we assessed how targeted gene deletion perturbed the model’s internal representation space. For each *in silico* knockout, we quantified the embedding drift (using cosine similarity and Euclidean distance) between the original and perturbed cellular states. To determine if this drift propagated according to true biological network topology, we classified the cellular gene space into three hierarchical tiers: direct interaction partners of the target gene (curated via SFARI), indirectly related ASD risk genes, and stably expressed control genes. By comparing the perturbation response across these tiers, we evaluated the model’s ability to natively reflect the cascading impact of genetic disruption across known regulatory networks. Consistent with true biological topology, we observed that both direct interactors and indirectly related ASD risk genes exhibited fundamentally greater embedding shifts compared to control genes (Fig. 3l).

### Embedding-derived functional hierarchy

To test whether DigitalBrain captures brain organization beyond a flat representation of dissociated cell states, we compared region-level hierarchies derived from DigitalBrain CLS embeddings with those derived from raw transcriptomic features. We focused on sliced Wasserstein distance (SWD), which compares full regional single-cell distributions without explicit cell-type matching, as a stringent test of whether the learned representation organizes anatomically defined regions in a biologically meaningful manner.

Both raw-SWD and embedding-SWD (Figure Table 2) produced non-flat and anatomically interpretable dendrograms, indicating that hierarchical regional structure is present in both representation spaces. At the level of local clusters, both methods recovered several biologically plausible modules. These included a frontoparietal control-like cluster linking posterodorsal (superior) parietal cortex, rostroventral portion of DFC (area 46) and ventrolateral prefrontal cortex, a visual extrastriate cluster linking area prostriata and parastriate cortex (area V2, area 18), and a core striatal cluster linking body of caudate, putamen and nucleus accumbens. Thus, the presence of compact functional or systems-like modules was not unique to the learned embedding.

**Table 2.**
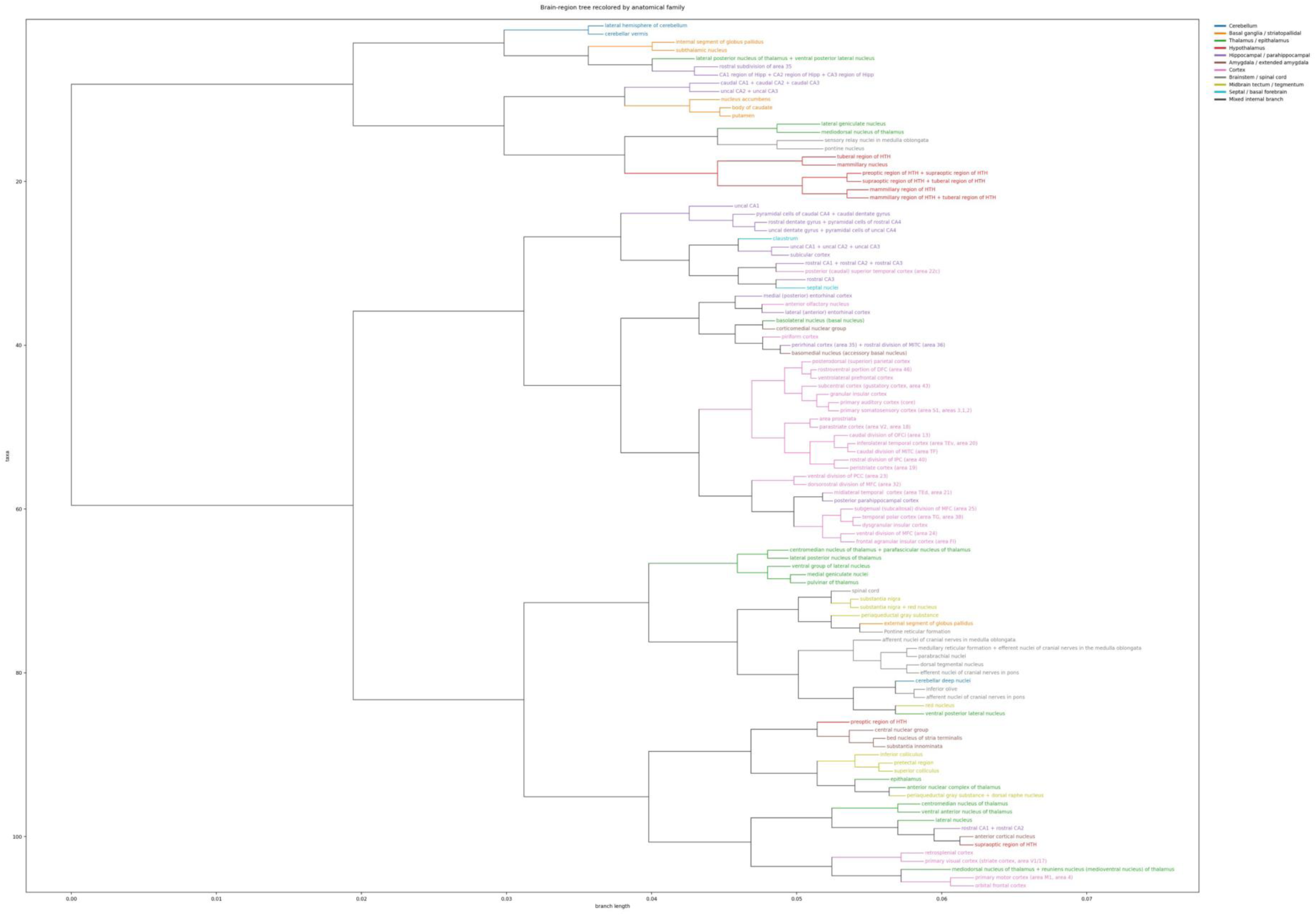
Embedding-derived hierarchy reveals higher-order organization of human brain regions. Region-level clustering based on DigitalBrain cls embeddings recovers biologically plausible local modules and more clearly organizes anatomically distinct regions into higher-order branches consistent with known distributed brain systems. This suggests that the learned embedding captures functional hierarchy beyond gross anatomical grouping.

The main difference emerged in how these local clusters were assembled into higher-order branches. In the embedding-SWD tree, several mixed-anatomy modules remained embedded within biologically coherent parent branches. For example, bed nucleus of stria terminalis and substantia innominata were joined by central nuclear group and then preoptic region of HTH, forming a hierarchical extended-amygdala–hypothalamic–basal forebrain branch consistent with a distributed autonomic-arousal system. Similarly, the striatal cluster (body of caudate, putamen, nucleus accumbens) was placed together with hippocampal fields (uncal CA2 + uncal CA3, caudal CA1 + caudal CA2 + caudal CA3), yielding a hippocampo-striatal higher-order branch consistent with contextual and action-related organization. The embedding tree also separated thalamic structure into more differentiated network-like branches, including mediodorsal nucleus of thalamus + reuniens nucleus grouped with orbital frontal cortex and primary motor cortex, rather than retaining all thalamic nuclei within a single block.

By contrast, in the raw-SWD tree, biologically plausible local modules were more often absorbed into less differentiated or harder-to-interpret super-clades. For example, although raw-SWD also recovered a septo-hippocampal branch linking rostral CA1 + rostral CA2 with septal nuclei, and an olfactory-related branch linking lateral (anterior) entorhinal cortex with piriform cortex, higher-order organization was less clearly resolved in several systems. Most notably, raw-SWD grouped a broad set of thalamic nuclei into a single large thalamic block, rather than distributing them into distinct cortical- or system-related branches. Raw-SWD also produced some less readily interpretable higher-order joins, including grouping primary visual cortex (striate cortex, area V1/17) with hippocampal CA-field branches, and grouping CA1 region of Hipp + CA2 region of Hipp + CA3 region of Hipp with external segment of globus pallidus.

Importantly, ambiguous cross-anatomical branches were not unique to the raw tree. The embedding-SWD tree also contained mixed branches that were difficult to interpret purely in systems terms, including a clade in which rostral subdivision of area 35 and CA1 region of Hipp + CA2 region of Hipp + CA3 region of Hipp were grouped within a broader branch containing internal segment of globus pallidus, subthalamic nucleus, and lateral posterior nucleus of thalamus + ventral posterior lateral nucleus. Thus, the distinction between the two methods is not that one preserves anatomy whereas the other does not, but that they differ in how local modules are organized into higher-order structure.

Taken together, these observations support a cautious but meaningful conclusion. Both raw-SWD and embedding-SWD recover biologically plausible local modules, but embedding-SWD more clearly organizes several of these modules into higher-order branches that are compatible with known distributed systems, particularly in thalamic, striatal, and extended-amygdala–basal forebrain organization. The comparison remains qualitative and is based on the biological interpretability of tree topology rather than on a formal statistical score. Accordingly, the observed branches should not be interpreted as direct reconstructions of anatomical connectivity or causal circuitry. Rather, the results suggest that the learned embedding captures a layer of functional hierarchy beyond gross anatomy by organizing anatomically distinct regions into transcriptomic modules and parent branches that are more consistent with distributed brain systems.

### DigitalBrain identifies multiscale aging-associated vulnerability in human dentate granule cells

Having established that DigitalBrain learns biologically meaningful gene representations, we next asked whether the model could support concrete biological discovery beyond standard integration and annotation benchmarks. As a proof-of-principle application, we applied DigitalBrain to cell type- and region-specific aging analysis in the human hippocampus. This provided a stringent test of whether the learned embedding space could recover interpretable and transferable aging-associated structure across multiple levels of analysis, from global gene regulatory network (GRN) organization to functional modules, pathway rewiring, and aging sensitive genes. We focused on the dentate gyrus because of its central role in hippocampal function, its importance for pattern separation, and its known sensitivity to age-related decline.

DGCs emerged as a particularly age-sensitive hippocampal cell population in the embedding analysis. In human hippocampal single-cell datasets^24,29^ (hereafter referred to as the Su dataset and the Zhou dataset), DGC embeddings from Su dataset (Fig. 4a and b) formed a distinct manifold with a clear age-associated gradient across 18 donors aged 0–92 years (Fig. 4c), suggesting that aging aligns with a major axis of DGC variation. We therefore used DGCs as a test case, with Su dataset as the discovery cohort and an independent DGC cells from Zhou dataset (28 donors, ages 0.6–92 years) for validation.

**Figure 4.**
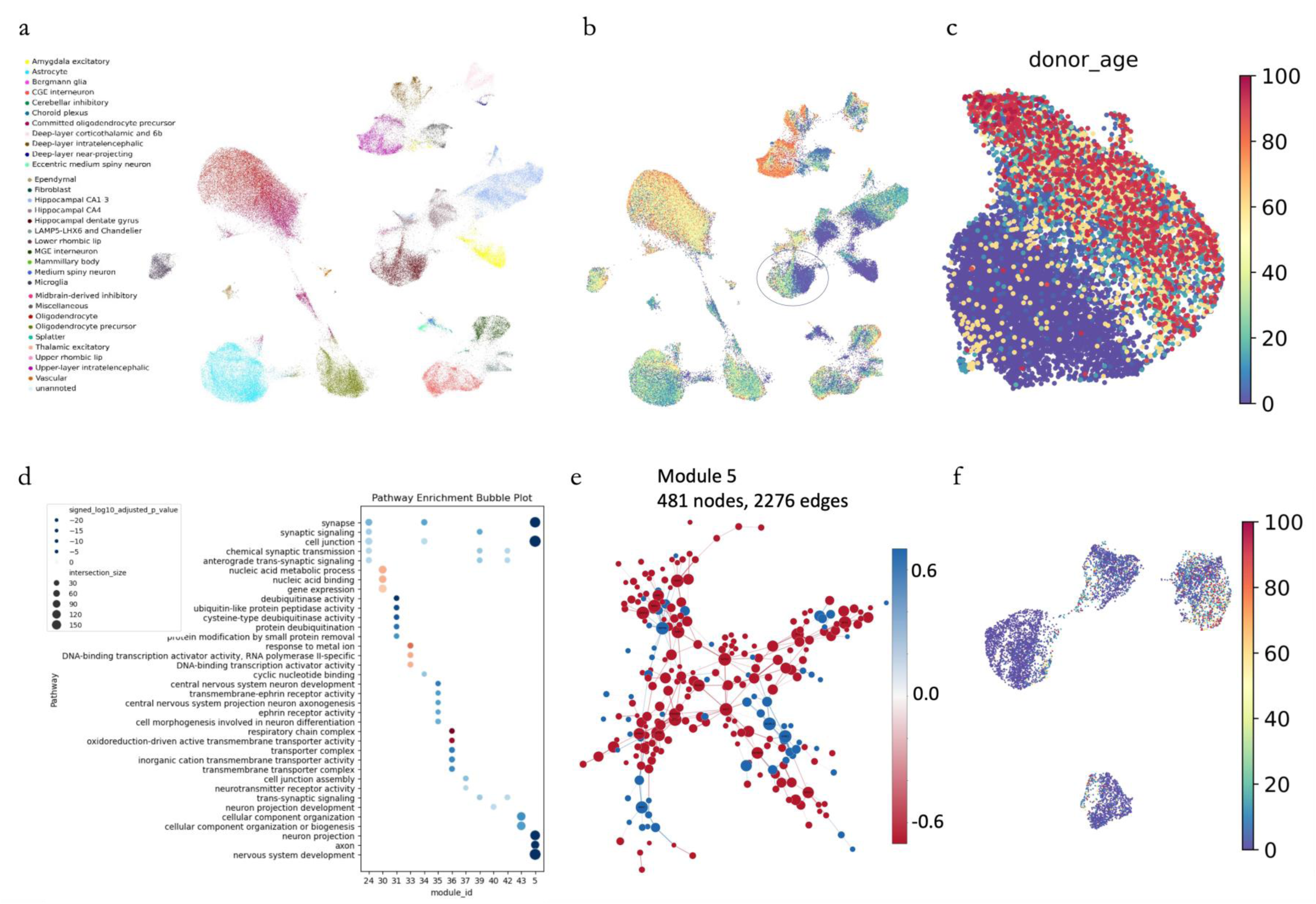
Cross-dataset aging analysis reveals conserved dentate gyrus network vulnerability across the lifespan. **(a)** UMAP of cell embeddings from the Su dataset. human hippocampal dataset after quality control (donors aged 0–92 years), coloured by annotated cell type. The dashed circle highlights dentate gyrus granule cells (DGCs), which were selected for downstream aging analysis. **(b)** The same embedding colored by donor age. Hexagons represent local neighborhoods of cells, with color indicating mean donor age after nearest-neighbor smoothing, revealing broad age-associated structure in the embedding. **(c)** UMAP of DGCs coloured by donor age, showing a clear age-associated gradient across the manifold, consistent with aging as a major continuous axis of variation in DGC state space. Red outlines indicate younger-enriched and older-enriched regions. **(d)** Functional enrichment of Leiden modules identified from the consensus mKNN+ network constructed in DigitalBrain gene-embedding space for DGCs. For each module, the top five significant go terms are shown (adjusted P < 0.05). Bubble size indicates overlap size and colour indicates signed −log10(adjusted P), with red denoting compensatory modules (intra-module connectivity increasing with age) and blue denoting declining modules (decreasing connectivity with age). **(e)** GRN of module 5 from (f). **(f)** Split distribution of *ASIC2* in UMAP.

In the Su dataset, donor-resolved network analysis showed a directional aging signal at the GRN level: 56.8% of testable consensus edges declined with age, and this pattern became stronger under stricter high-confidence edge filters (p = 0.002). In contrast, the Zhou dataset showed no clear global directional signal, with 50.3% of edges declining with age (p = 0.480) and no significant bias in module direction (p = 0.972). These results indicate that broad network-wide connectivity loss is not uniformly reproducible across cohorts, prompting us to ask whether aging-related convergence emerges more clearly at finer levels of organization.

At the module level, aging-associated changes in the Su dataset were concentrated in neuronal and synaptic systems. Among 51 modules, 4 showed significant connectivity decline, and most age-associated modules trended downward (Fig. 4d). In the Zhou dataset, 11 of 100 modules showed significant changes, including 9 declining modules, again pointing to neuronal, axonal, and synaptic functions. These aging-associated modules were not simply the largest in the network, arguing against a size-driven effect. Other functionally coherent modules, including those related to transcriptional regulation, proteostasis, transport, and respiratory-chain activity, were detected but did not show significant aging association. Overall, the cross-dataset comparison consistently highlighted synaptic and neuronal systems as the modules most affected by aging, in both broad networks and smaller, more specific subnetworks.

A stricter pathway-level analysis in the Su dataset further highlighted declining neuronal and synaptic connectivity, including pathways related to postsynaptic structure, synaptic transmission, NMDA receptor signaling, long-term potentiation, axon guidance, and glutamatergic function. No pathways reached significance in the Zhou dataset under the same criterion. Thus, pathway-level significance was strongest in the discovery cohort, whereas cross-dataset convergence was more evident at the level of broader functional modules.

At gene level, cross-dataset analysis revealed strong convergence of aging sensitive genes despite weaker modular consistence in the Zhou dataset. The Su and Zhou datasets yielded partially overlapping but highly concordant gene sets, with 14 shared genes among the aging sensitive genes (hypergeometric test for overlap, *P* = 6.53 × 10^14^). These recurrent genes: *ASIC2* (Fig. 4e)*, CLSTN2, DAB1, RFX3, KHDRBS2, FTX, KCNQ5, PTPRG, DNM3, PROX1, PRKG1, REV3L, GRIP1,* and *PITPNC1*, converged on related neuronal functions, including synaptic organization, membrane excitability, axon guidance/connectivity, and neuronal regulatory identity. This functional consistency was supported by independent analyses. Gene set enrichment of the ranked gene lists by GSEA^30^ highlighted shared neural-related terms across datasets, particularly voltage-gated ion channel activity and synaptic functions (for Zhou dataset, six terms at FDR < 0.05, including voltage-gated monoatomic ion channel activity at FDR = 0.021 and NGF-stimulated transcription at FDR = 0.045; for Su dataset, near-significant trend for these exact shared pathways with minimum FDR = 0.051). Top-ranked aging sensitive genes also showed strong local consistency: most were embedded in neighborhoods that likewise exhibited age-associated decline, indicating that these genes mark focal points of local network deterioration rather than isolated gene-level changes. In addition, network-based decline scores were positively correlated with expression-based age effects in both datasets, showing broad agreement between connectivity loss and transcriptomic aging trends.

As an illustrative example, *ASIC2* embeddings showed a structured embedding manifold with four separable clusters (Fig. 4f) and a non-random distribution of donor age, with young and old donors enriched in different clusters. This pattern suggests that the embedding captures heterogeneous, age-associated contexts for the same gene across the lifespan. Notably, this gene-level structure is consistent with the broader age-associated heterogeneity observed in DGCs, indicating that aging-related variation is reflected at both the cellular and gene-embedding levels.

Overall, the consistency of cross-dataset aging signals was scale-dependent. Both datasets converged on related neuronal and synaptic modules and showed significant overlap in aging sensitive genes. This pattern may indicate that, in heterogeneous human donor datasets, reproducibility is strongest at the level of functional modules and aging sensitive genes rather than in uniform global network measures. Together, these analyses show that DigitalBrain can move beyond benchmark performance to support biologically interpretable, multiscale discovery in human aging data.

### Age-associated structure is captured in both cell and gene embeddings in the healthy hippocampus

The GRN-based analysis above focused on age-dependent rewiring of relationships among genes. At the cell level, we then measure the donor-specific GRN changed with age within each cell type, which reveals that age-associated GRN drift was strongest in astrocytes and dentate gyrus granule cells (DGCs), with additional signal in CA1-3 pyramidal neurons and oligodendrocyte precursor cells (Supplementary Fig. 3).

The gene-embedding manifolds (Fig. 4f) suggested that aging might also be reflected in the geometry and distribution of the embedding space itself. We therefor developed an SWD-based analysis to quantify age-associated drift in cell and gene embeddings. In this analysis, greater embedding drift indicates stronger age-associated reorganization in the learned representation space, and can therefore be used to identify cell populations and genes that may be sensitive to aging. Based on our current results, the top 5 ranking genes are highly heterogeneous across different cell types (Fig. 5a and b), indicating that aging-related gene vulnerability is cell-type-specific rather than homogeneous. Both high and low-variability genes, such as *NRXN1* and *MALAT1*, showed marked cell type dependent distribution in gene-embedding space (Fig. 5c and d). Together, these results identify cell types and genes with stronger age-associated embedding drift as prioritized candidates for further functional investigation.

**Fig. 5.**
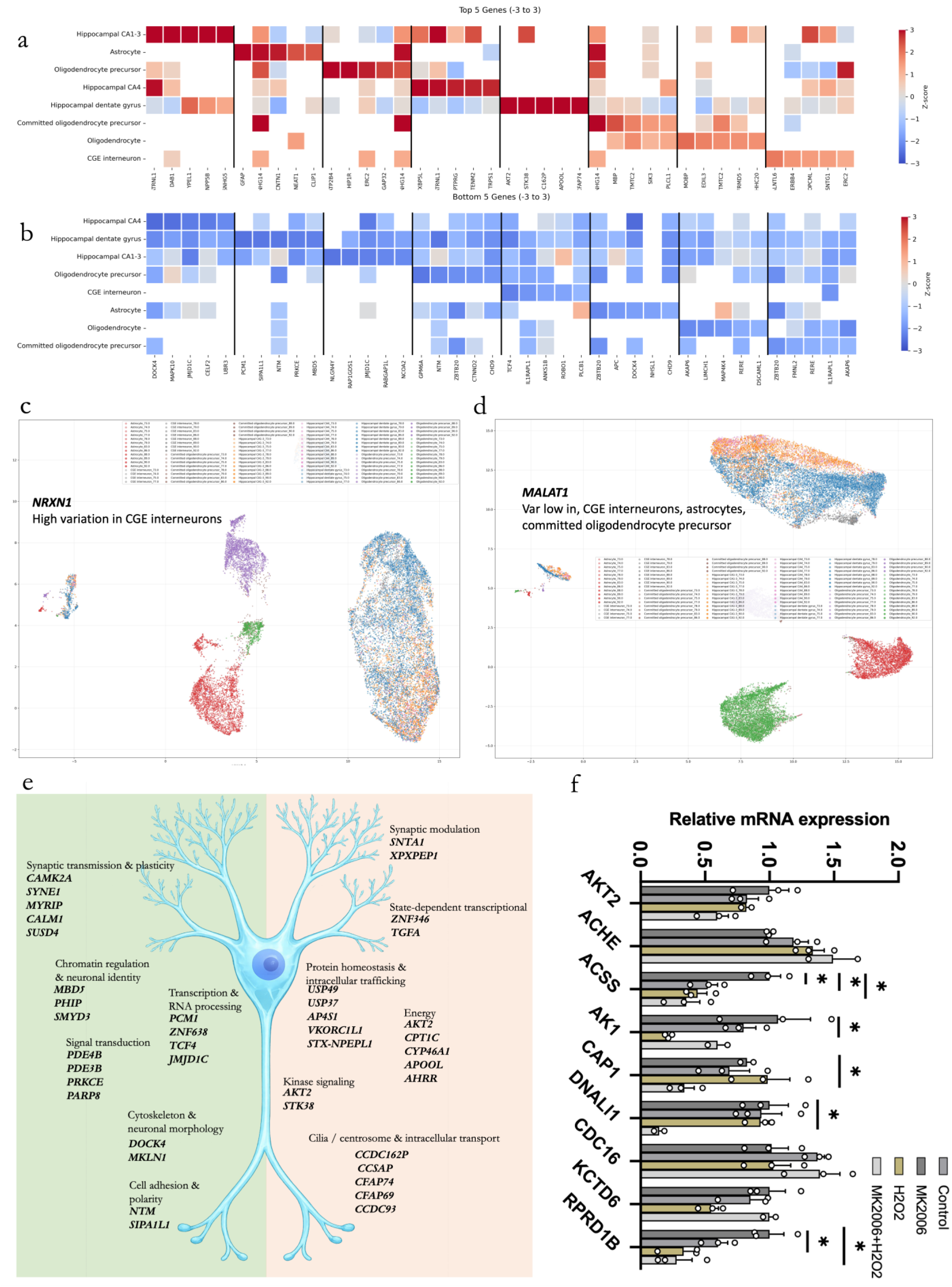
Age-associated structure in DigitalBrain cell and gene embeddings in the healthy hippocampus. **(a)** Heatmap of the top five genes with the highest age-variability scores in each cell type. Scores were normalized within cell type as Z-scores relative to a housekeeping-gene baseline. Rows represent cell types and columns represent selected genes grouped by source cell type; warmer colors indicate higher age-related variability. **(b)** Heatmap of the top five genes with the lowest age-variability scores in each cell type. **(c)** UMAP visualization of representative high-variability genes in gene-embedding space, illustrated here for *NRXN1*. **(d)** UMAP visualization of representative low-variability genes in gene-embedding space, illustrated here for *MALAT1*. **(e)** Gene state dynamics in human dentate granule cells (DGCs) across late-life aging (70–93 years), analyzed using embedding-based functional representations. Genes were ranked by embedding variance across age bins to distinguish functionally stable genes from dynamically reconfiguring genes. Neuronal identity, synaptic, chromatin, and structural genes remain embedded in stable regions of the functional gene manifold, whereas genes involved in metabolism, energy regulation, proteostasis, and intracellular trafficking exhibit progressive, age-dependent gene state drift. Peripheral network views illustrate age-specific reorganization of gene embedding neighborhoods across successive age bins. These patterns emerge directly from learned representations without prior biological annotation, revealing structured, programmed functional reconfiguration during human brain aging. **(f)** Relative mRNA expression levels of *AKT2* and model predicted *AKT2* associated genes (*ACHE, ACSS, AK1, CAP1, DNALI1, CDC16, KCTD6, RPRD1B*); n = 3. Two-way ANOVA; Statistical analysis was performed for each gene vs Control. \**p <* 0.05, \*\**p <* 0.01, \*\*\**p <* 0.001. All data are presented as mean ± SEM.

Because DGCs showed one of the strongest age-associated signals, we examined gene-embedding dynamics in this population during late-life aging (70–93 years). Ranking genes by embedding drift across age bins separated relatively stable gene programs from those showing progressive age-associated drift (Fig. 5e). Based on their annotated functions in the literature, we grouped these genes into broad categories including neuronal identity, synaptic, chromatin, and structural functions, whereas genes with greater drift were associated with metabolism, energy regulation, proteostasis, and intracellular trafficking. To test whether embedding proximity reflects functional relatedness, we perturbed *AKT2*, a top-ranked gene by age-associated embedding drift with a well-defined local neighborhood in embedding space. Pharmacological inhibition of AKT2 activity with MK2206 and H_2_O_2_ induced oxidative stress, the latter used here as a senescence-like aging model in primary hippocampal neurons, altered the expression of several model-predicted *AKT2* associated genes, including *ACHE*, *ACSS*, *AK1*, *CAP1*, *KCTD6* and *RPRD1B*, providing preliminary experimental support that local neighborhoods in embedding space capture biologically meaningful relationships (Fig. 5f). This supports the biological relevance of the embedding-defined local network and suggests that nearby genes can share related responses to perturbation.

## Discussion

In this work, we developed DigitalBrain, a human brain-specific framework that combines a harmonized whole-brain single-cell atlas DigitalBrain-Atlas, a representation model DigitalBrain-M1 trained with human brain data and embedding based analysis methods. DigitalBrain-Atlas contains 109 datasets comprising more than 16 million cells across 165 brain regions, the whole lifespan and multiple disease contexts, representing a structured and valuable resource for modeling human brain cellular diversity. We then used this to train DigitalBrain-M1 to learn coupled cell and gene representations and showed that it supports integration, annotation and interpretable analysis across datasets and biological scales. Applying this framework to hippocampal aging highlighted dentate gyrus granule cells and associated gene programs as prominent features of aging-related state change. Importantly, cross-dataset analysis further showed that the model recovers consistent age-associated modules and recurrent driver genes in drifting GRN across independent DGC datasets, supporting the generalizability and robustness of the learned representation.

More broadly, our work can be viewed as an extension of recent efforts toward a virtual cell, in which large-scale transcriptomic models learn computational representations of cellular identity and state^14–18^. These studies have shown that learned representations can capture biologically meaningful structure and support integration, annotation and perturbation-oriented analysis. However, cells in vivo are not isolated entities: they are embedded within organs that are organized across anatomical, developmental and pathological axes. For the human brain in particular, cellular states are shaped by regional context, lifespan stage and disease condition in addition to intrinsic cell identity. Our results suggest that brain-specific modeling benefits from explicitly learning within this broader organ-level context.

From this perspective, a minimal form of a virtual organ should include at least two components. First, it requires a sufficiently broad and hierarchically organized atlas that samples the major anatomical regions and cellular populations of the organ across relevant biological conditions. Second, it requires a representation framework that can learn from this structured resource while preserving information across multiple biological scales. DigitalBrain is an initial step in that direction. DigitalBrain-Atlas brings together more than 16 million cells across 165 brain regions, the human lifespan and multiple clinical conditions within a harmonized anatomical framework, and DigitalBrain-M1 is trained on this structured resource rather than on a relatively flat collection of dissociated cells. We therefore view DigitalBrain not as a complete virtual organ, but as an early brain-specific framework for representing cellular states within a broader organ-level hierarchy. We therefore view DigitalBrain not as a complete virtual organ, but as a first step toward a brain-specific digital organ framework that links cellular representations to organ-level structure.

The results suggest that this framework captures biologically meaningful structure at several levels. At the cell level, the model supports integration and annotation while preserving major biological organization. At the gene level, the learned embeddings reflect cell-type specificity, transcriptional concordance and pathway-level coherence, and support perturbation-oriented analyses of disease-relevant genes. At the regional level, embedding-based SWD recovered mixed-anatomy modules that, in several cases, aligned with known distributed systems more readily than analogous structures in raw transcriptomic space. Together, these findings suggest that DigitalBrain begins to connect molecular, cellular and regional organization within a shared computational framework.

The significance of this framework is therefore not that it produces hierarchy per se, but that it provides a tractable way to organize fragmented whole-brain single-cell data into a biologically interpretable representation across scales. In the present version, DigitalBrain addresses a foundational layer of the digital brain problem: the representation of whole-brain cellular and molecular organization from transcriptomic data. This creates a scaffold on which richer and more comprehensive models can be built.

A natural next stage is to extend this framework from molecular and cellular organization toward brain dynamics. For the brain, a fuller digital-organ representation will likely require integrating transcriptomic state with spatial architecture, physiological activity, neural rhythms, imaging-derived organization and longitudinal state transitions across aging and disease. Future development could therefore combine DigitalBrain-like embeddings with spatial transcriptomics, electrophysiology, perturbation datasets and multimodal brain imaging, enabling models that connect molecular state to functional dynamics across scales. In this view, the current study provides an initial molecular and cellular scaffold for a more complete digital brain.

More broadly, our results suggest that progress toward a digital organ will depend on three elements: a sufficiently broad and harmonized atlas, an organ-specific representation model and analyses that connect learned structure back to interpretable biology. DigitalBrain brings these components together for the human brain and establishes a practical framework for studying how molecular states relate to anatomical hierarchy, aging and disease. Although still an early version, it represents a concrete step from the virtual cell toward a digital brain model that is increasingly multiscale, dynamic and biologically grounded.

---

**Supplementary Figure 1.**
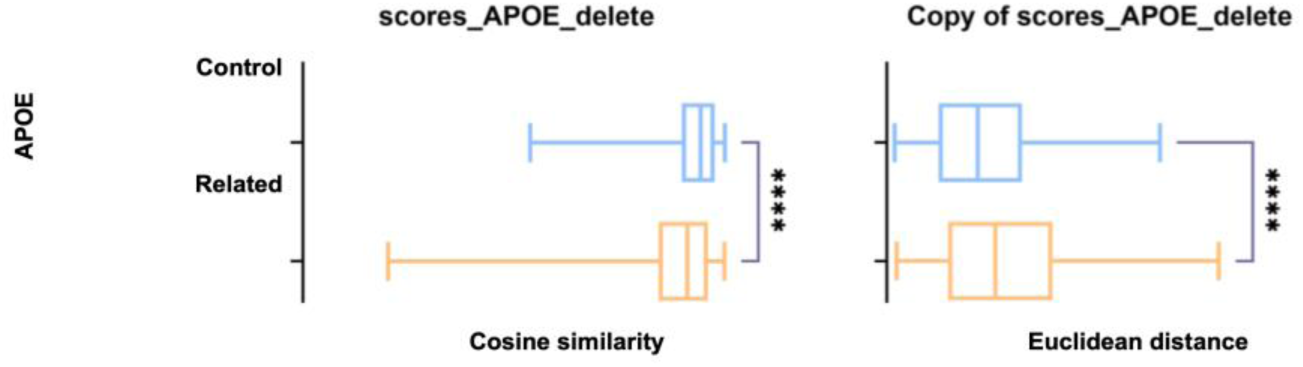
Evaluation of embedding variations before and after *APOE* knockout. Boxplots display the Cosine Similarity and Euclidean Distance for the Control versus Related gene groups. Statistical significance is indicated by asterisks (**** P < 0.0001, ** P < 0.01, * P < 0.05, ns: not significant). The center line in the boxplots represents the median, box limits indicate the upper and lower quartiles, and whiskers represent the 1.5× interquartile range.

**Supplementary Figure 2.**
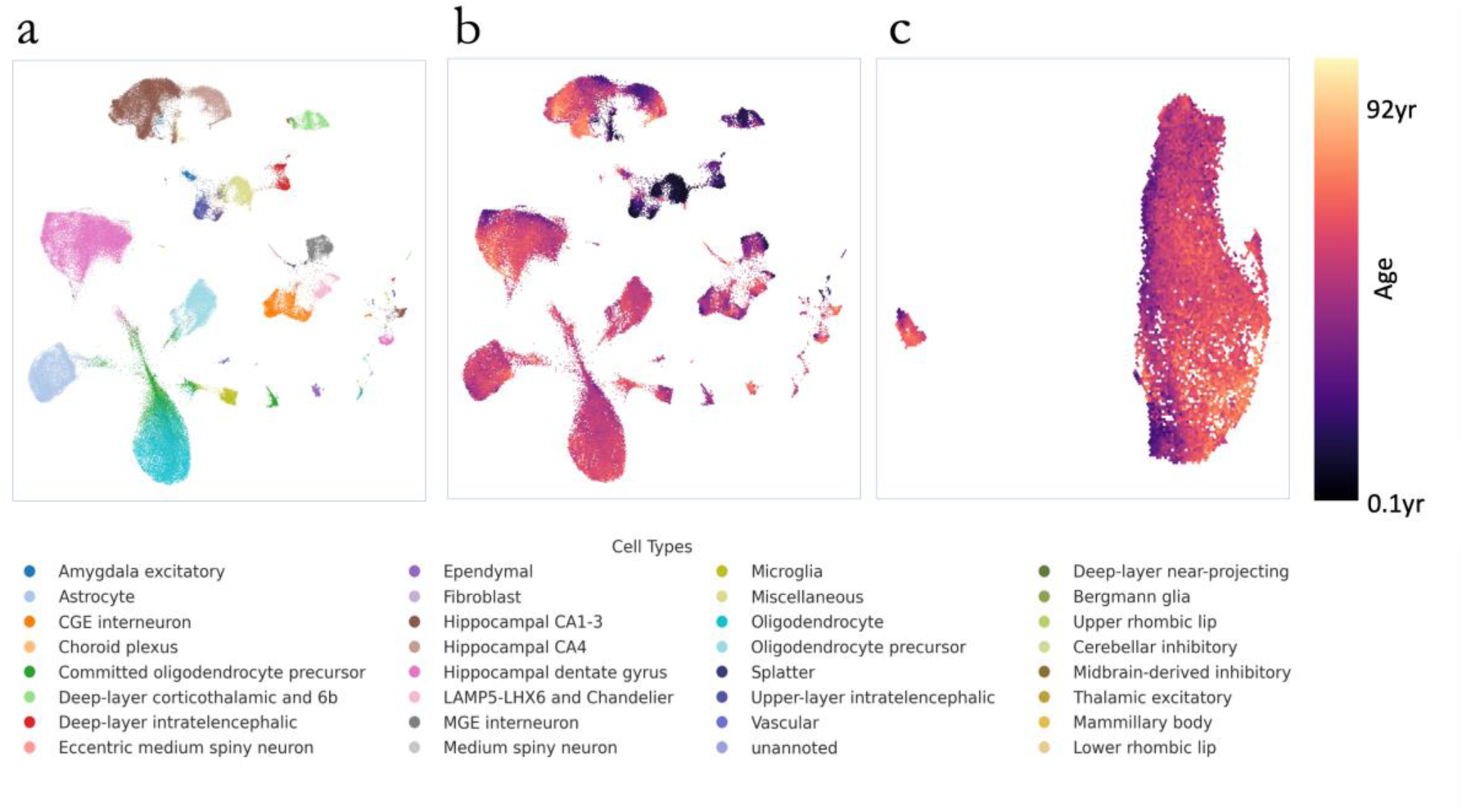
(a) Global UMAP of hippocampal single-cell embeddings, colored by major cell type, showing separation of major populations. **(b)** The same embedding colored by donor age. Hexagons represent local neighborhoods of cells, with color indicating mean donor age after nearest-neighbor smoothing, revealing broad age-associated structure in the embedding. **(c)** Age-colored UMAP of dentate gyrus granule cells.

**Supplementary Figure 3.**
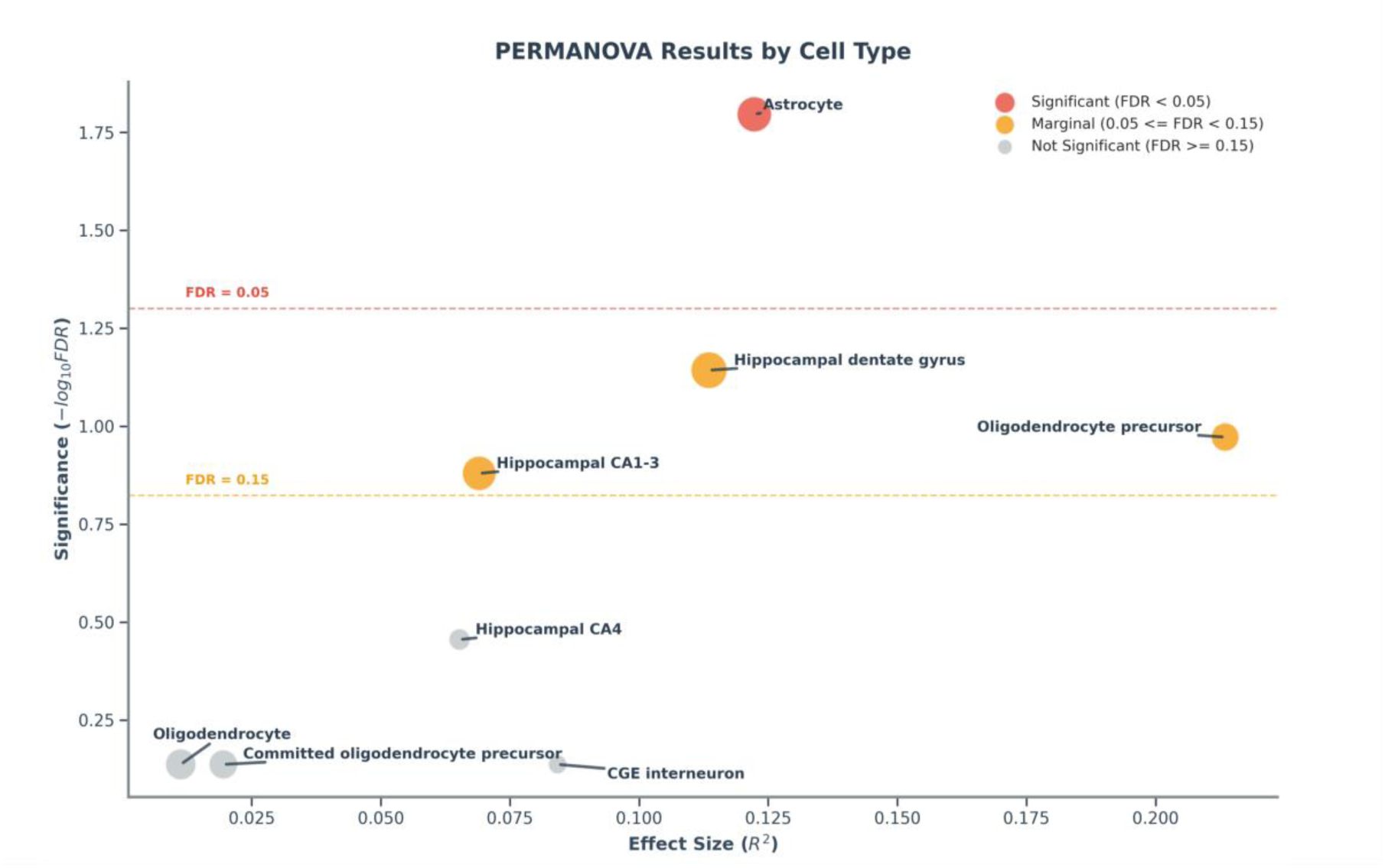
PERMANOVA-based summary of age effects across cell types. The x axis shows effect size (R ²), and the y axis shows significance as −log10(FDR). Dashed lines indicate FDR thresholds of 0.15 and 0.05. Astrocytes and dentate gyrus granule cells show the strongest age-associated effects, with additional signal in CA1–3 pyramidal neurons and oligodendrocyte precursor cells.

